# A single cell transcriptome atlas of the developing zebrafish hindbrain

**DOI:** 10.1101/745141

**Authors:** Monica Tambalo, Richard Mitter, David G. Wilkinson

## Abstract

Segmentation of the vertebrate hindbrain leads to the formation of rhombomeres, each with a distinct anteroposterior identity. Specialised boundary cells form at segment borders that act as a source or regulator of neuronal differentiation. In zebrafish, there is spatial patterning of neurogenesis in which non-neurogenic zones form at bounderies and segment centres, in part mediated by Fgf20 signaling. To further understand the control of neurogenesis, we have carried out single cell RNA sequencing of the zebrafish hindbrain at three different stages of patterning. Analyses of the data reveal known and novel markers of distinct hindbrain segments, of cell types along the dorsoventral axis, and of the transition of progenitors to neuronal differentiation. We find major shifts in the transcriptome of progenitors and of differentiating cells between the different stages analysed. Supervised clustering with markers of boundary cells and segment centres, together with RNA-seq analysis of Fgf-regulated genes, has revealed new candidate regulators of cell differentiation in the hindbrain. These data provide a valuable resource for functional investigations of the patterning of neurogenesis and the transition of progenitors to neuronal differentiation.

## Introduction

Development of the central nervous system (CNS) requires precise regulation of the differentiation of neuronal and glial cell types from neural progenitor cells. This is achieved through a network of cell-cell signaling and transcription factors that inhibit or promote cell differentiation and specify cell type along the dorsoventral and anteroposterior axes of the neural epithelium. Cell specification along the dorsoventral axis involves localised sources of Shh, BMP and Wnt signals that act in a concentration-dependent manner to regulate expression of specific transcription factors (Dessaud et al., 2008; Dessaud et al., 2007; Hikasa and Sokol, 2013; Ikeya et al., 1997; Lee and Jessell, 1999; Liem et al., 1997; Panhuysen et al., 2004; Timmer et al., 2002; Ulloa and Marti, 2010). This positional information is integrated with patterning along the anteroposterior axis, which regulates expression of transcription factors that specify regional identity within the brain and spinal cord (Alexander et al., 2009). Differentiation is also under temporal regulation, with distinct neuronal or glial cell types arising at different times (Guillemot, 2007). It is essential that a pool of progenitor cells is maintained as a source of later-differentiating cells, and this is achieved by multiple mechanisms that inhibit differentiation.

The switch of progenitor cells to neuronal differentiation involves sustained high-level expression of proneural transcription factors that initiate a cascade of gene expression leading to expression of terminal neuronal markers (Bertrand et al., 2002). The expression and function of proneural genes is antagonised by intrinsic factors, as well as by extrinsic signals such as Notch ligands and Fgfs that inhibit differentiation (Fisher and Caudy, 1998; Gonzalez-Quevedo et al., 2010; Kageyama et al., 2005; Ortega et al., 1998; Vaccarino et al., 1999; Zheng et al., 2004). In some regions of the developing CNS, neurogenesis occurs widely, and Notch-mediated lateral inhibition ensures that progenitor cells are maintained (Pierfelice et al., 2011). In other regions, neurogenesis is patterned by spatially-resticted expression of Hes/Her genes that inhibit neuronal differentiation (Bae et al., 2005; Geling et al., 2003). Studies of the vertebrate hindbrain have revealed further mechanisms that pattern neuronal differentiation.

At early stages, the neural epithelium of the hindbrain is subdivided to form seven rhombomeres (r1-r7), each expressing a distinct set of transcription factors, including krox20, mafB, vhnf1 and hox genes, that underlie segmentation and anteroposterior identity (Alexander et al., 2009). A similar but different set of neurons is generated in each rhombomere (Clarke and Lumsden, 1993; Lumsden, 2004; Lumsden and Keynes, 1989). There is a partial understanding of mechanisms that link A-P identity to neuronal cell type specification in the hindbrain (Narita and Rijli, 2009).

Boundary formation has a crucial role in the organisation of neurogenesis in the hindbrain. Through cell identity regulation (Addison et al., 2018; Wang et al., 2017) and Eph-ephrin mediated cell segregation (Batlle and Wilkinson, 2012; Cayuso et al., 2015; Fagotto, 2014), each rhombomere is demarcated by sharp borders and has a homogeneous segmental identity. Specialised boundary cells form at each rhombomere border (Guthrie and Lumsden, 1991), which express specific molecular markers (Cheng et al., 2004; Cooke et al., 2005; Heyman et al., 1995; Letelier et al., 2018; Xu et al., 1995). These boundary cells are induced by Eph receptor signaling that leads to activation of Taz (Cayuso et al., 2019). In the chick hindbrain, boundary cells have a lower rate of proliferation (Guthrie et al., 1991) and are Sox2-expressing stem cells that are a source of neurogenesis (Peretz et al., 2016). A different situation occurs in the zebrafish hindbrain, in which expression of proneural transcription factors is initially widespread, and later becomes confined to zones flanking hindbrain boundary cells (Amoyel et al., 2005; Cheng et al., 2004). Notch activation promoted by *rfng* expression inhibits neurogenesis in boundary cells (Cheng et al., 2004). In addition, there is increased proliferation and inhibition of neurogenesis in boundary cells by activation of the Yap/Taz pathway downstream of mechanical tension, which declines at later stages (Voltes et al., 2019). Neurogenesis is inhibited at segment centres by *fgf20*-expressing neurons that act on the adjacent neural epithelium (Gonzalez-Quevedo et al., 2010) and are clustered by semaphorin-mediated chemorepulsion from boundary cells (Terriente et al., 2012). In addition to suppressing neuronal differentiation, Fgf signaling may switch progenitors at the segment centre to glial differentiation (Esain et al., 2010). The zebrafish hindbrain thus has a precise organisation of signaling sources that underlies a stereotyped pattern of neurogenic and non-neurogenic zones.

We set out to identify further potential regulators of neurogenesis during hindbrain segmentation by single cell RNA sequencing (scRNA-seq) to identify genes specifically expressed in distinct progenitors and differentiating cells, prior to and during the patterning of neurogenesis. Analyses of the transcriptome of single cells revealed known genes and new markers of distinct hindbrain segments, of cell types along the dorsoventral axis, and of the transition of progenitors to neuronal differentiation. We also find temporal changes in gene expression, both in progenitors and differentiating cells, at the different stages analysed. By carrying out supervised clustering, we have identified further genes specifically expressed in hindbrain boundary cells and segment centres. These findings are compared with bulk RNA-Seq analyses following loss and gain of Fgf signaling to identify potential regulators expressed in segment centres.

## RESULTS

### Single-cell profiling of the developing zebrafish hindbrain and surrounding tissues

To further understand the progressive patterning of neurogenesis in the developing zebrafish hindbrain, we analysed the transcriptome of single cells at three developmental stages (Fig.1A, B): 16 hpf (prior to patterning of neurogenesis), 24 hpf (beginning of neurogenic patterning) and 44 hpf (pattern of neurogenic and non-neurogenic zones fully established). For each stage, we micro-dissected the hindbrain territory from around 40 embryos, which were pooled. After enzymatic digestion and mechanical dissociation, the single-cell suspension was loaded into the droplet-based scRNA-seq platform 10X Genomics Chromium (Fig.1C). In total, 9026 cells were sequenced, with an average number of UMIs of 6916 and 1703 median genes per cell (Suppl. Fig.1). After applying the appropriate Seurat quality filters, a total of 2873, 2476 and 3497 cells - for 16 hpf, 24 hpf and 44 hpf embryos respectively - were retained for analysis.

**Figure 1.**
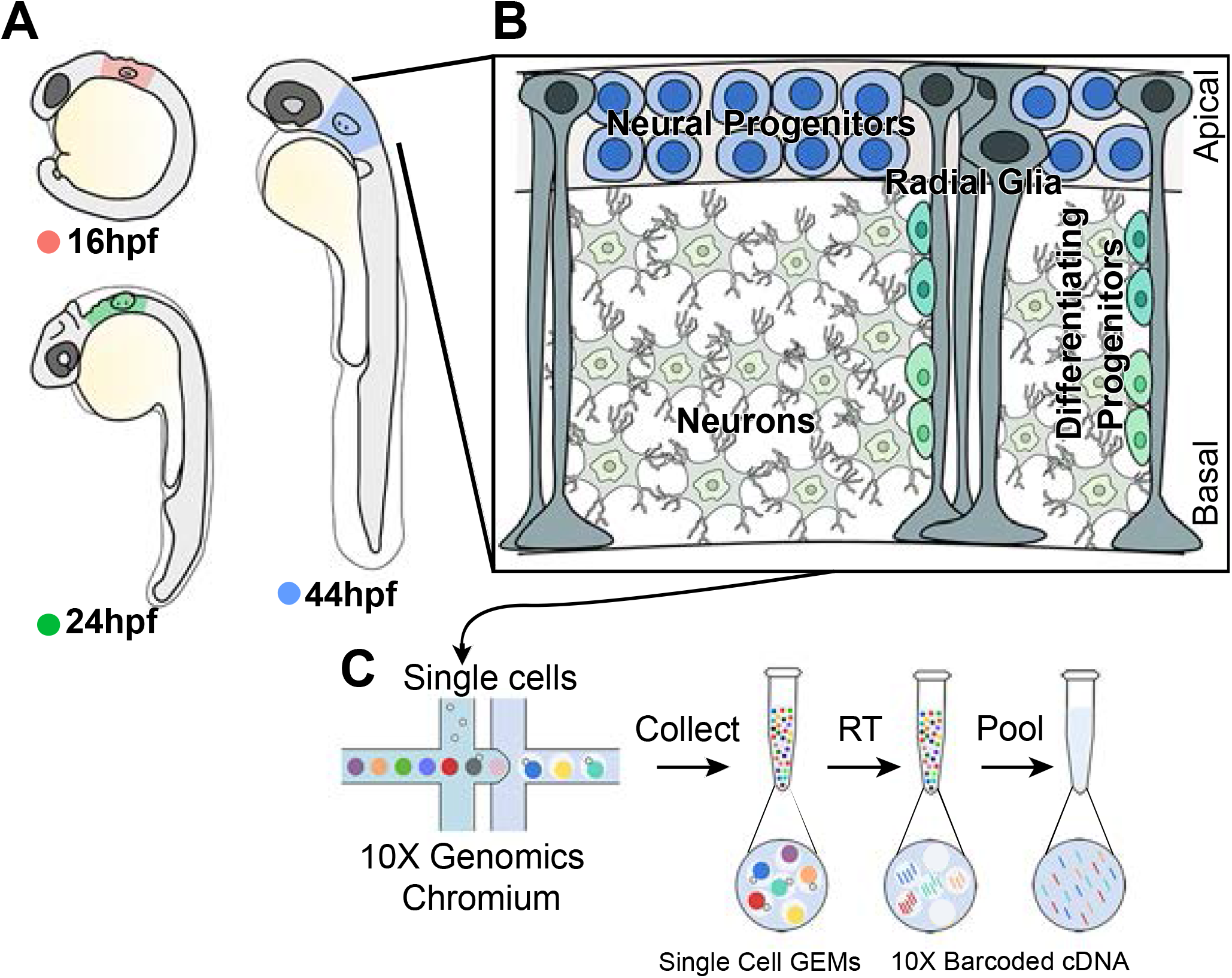
High throughput scRNA-seq strategy from the developing hindbrain. (A) The hindbrain of 16 hpf (pink), 24 hpf (green) and 44 hpf (blue) embryos was collected for scRNA-seq. (B) Drawing of the stereotypical hindbrain cell composition at 44 hpf. Progenitors and radial glia cell bodies occupy the ventricular region, while differentiating progenitors and neurons are in the mantle zone. (C) Schematic of the 10X Genomics Chromium workflow.

Seurat unsupervised clustering was used to classify cell population identity (Butler et al., 2018a; Stuart et al., 2018) for each stage and after aggregating the data from all stages (Suppl. Fig.2). Stage-specific cluster projection onto tSNE (t-distributed Stochastic Neighbour Embedding) plots revealed a central tight group of cells, with some substructure, surrounded by peripheral clusters (Suppl. Fig.2A-D). Since the dissections included tissues adjacent to the hindbrain, it is likely that the clusters correspond to distinct tissue types. We therefore used tissue marker genes to assign cluster identity. The progenitor marker Sox3 and neuronal gene Elavl4 have complementary expression, and together define the clusters derived from hindbrain territory (Suppl. Fig.2A’-D’, A”-D”). The main tissues found next to the hindbrain are the otic vesicle (Six1+, Neurod1+), cranial ganglia (Neurod1+), neural crest (Twist1+) and head mesenchyme (Colec12+). As expected, their expression is largely confined to the clusters surrounding the hindbrain domain (Suppl. Fig. 2A’-D’). Based on this analysis, we bioinformatically recovered hindbrain cells for each stage: 1821 cells at 16 hpf, 1600 cells at 24 hpf and 2719 cells at 44 hpf (Suppl. Table 1).

### Overall changes in hindbrain tissue composition

Using an unsupervised graph-based clustering approach we identified 9, 7 and 12 clusters at 16 hpf, 24 hpf and 44 hpf, respectively. Data sets were visualized with tSNE dimensionality reduction, and this revealed unique features that reflect the greatest transcriptomic differences between cell types at each developmental stage (Fig.2A, Fig. 3A, Fig.4A). Analysis of the top 30 significantly enriched genes per each cluster, and expression of known molecular markers, enabled us to assign identity to each cluster. The 16 hpf hindbrain is mainly constituted of progenitors (91% of total hindbrain cells; Fig.2A), and neurogenesis accounts for 6% of hindbrain cells. Progenitors remain the most abundant hindbrain cell type at 24 hpf (71% of hindbrain cells), while 28% of cells express markers of neuronal differentiation (Fig.3A). By 44 hpf, the proportion of progenitor cells has further diminished to 40%, with 55% of the cells expressing neuronal differentiation markers (Fig.4A). The clustering of cells by transcriptomic differences changes at the three stages. At 16 hpf, clustering is driven by segmental and dorsoventral identity (Fig.2A), whereas at 24 hpf and 44 hpf cells are clustered by dorsoventral identity and differentiation state (Fig. 3A, Fig.4A). This change reflects the greater proportion of cells undergoing differentiation at the later stages, with an increasing number of neuronal sub-types that are segregated into seven clusters by 44 hpf (Fig.4A). Below, we present more detailed analyses of each of these features that reveal known genes and novel markers of segmental identity, dorsoventral identity and differentiation state. An annotated list including information on any previous studies of these genes is presented in Supplementary Table 2.

**Figure 2.**
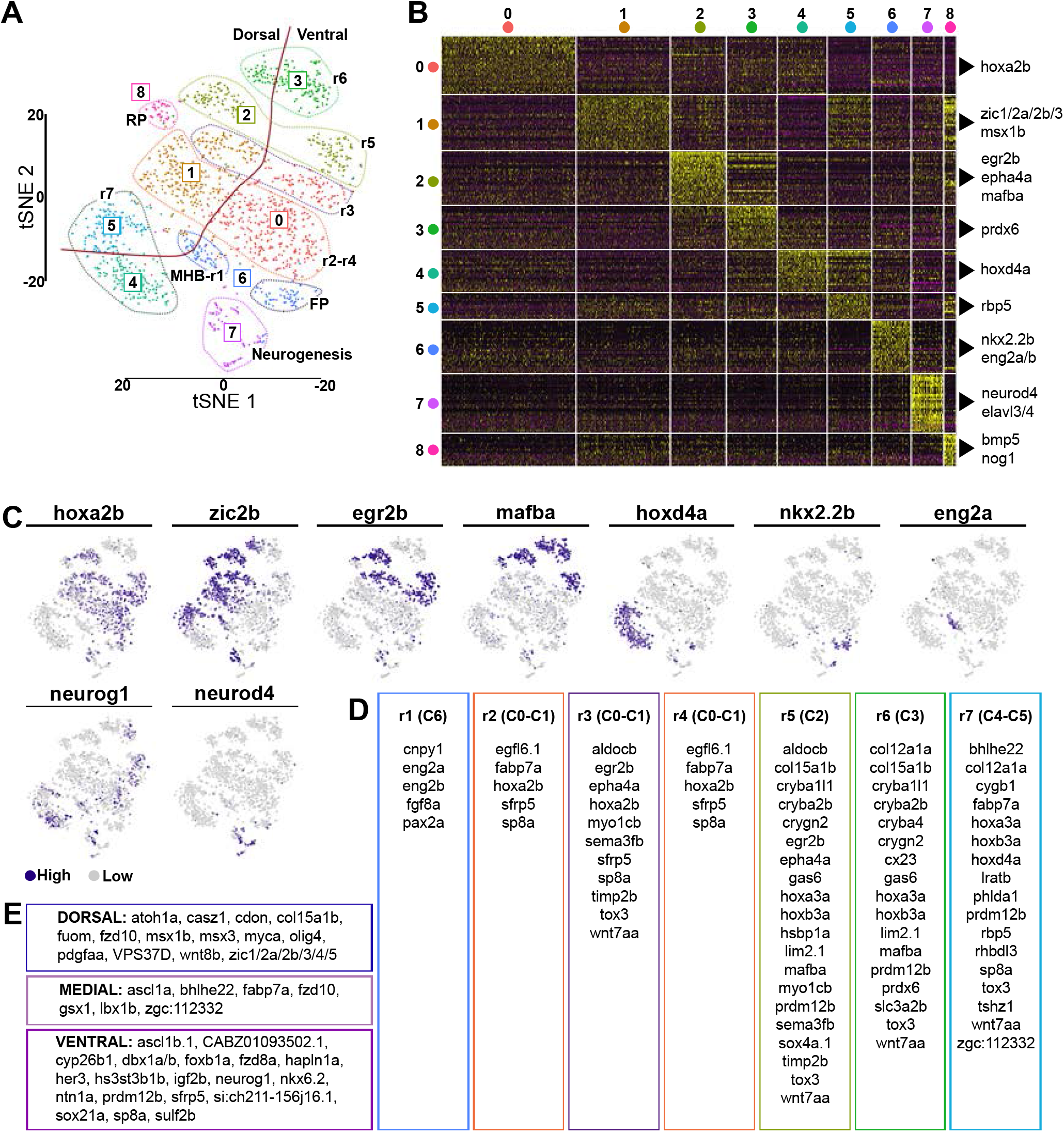
Cell population composition and signatures of the 16 hpf hindbrain. (A) Unsupervised tSNE plot subdivides hindbrain cells into 9 clusters (C0-C8). Dotted lines segregate different rhombomeres (r), midbrain-hindbrain boundary (MHB), floor plate (FP), roof plate (RP) and cells undergoing neurogenesis. The red line separates dorsal versus ventral cells. (B) Heatmap of the top 30 genes significantly enriched in each cluster; representative gene names are shown close to each cluster. The full gene list is in Supplementary File 1. (C) tSNE plots showing the log normalised counts of representative genes. Colour intensity is proportional to the expression level. (D) Summary of rhombomere-specific genes extracted from the top 30 significantly enriched. (E) Summary of genes restricted along the dorsoventral axis.

**Figure 3.**
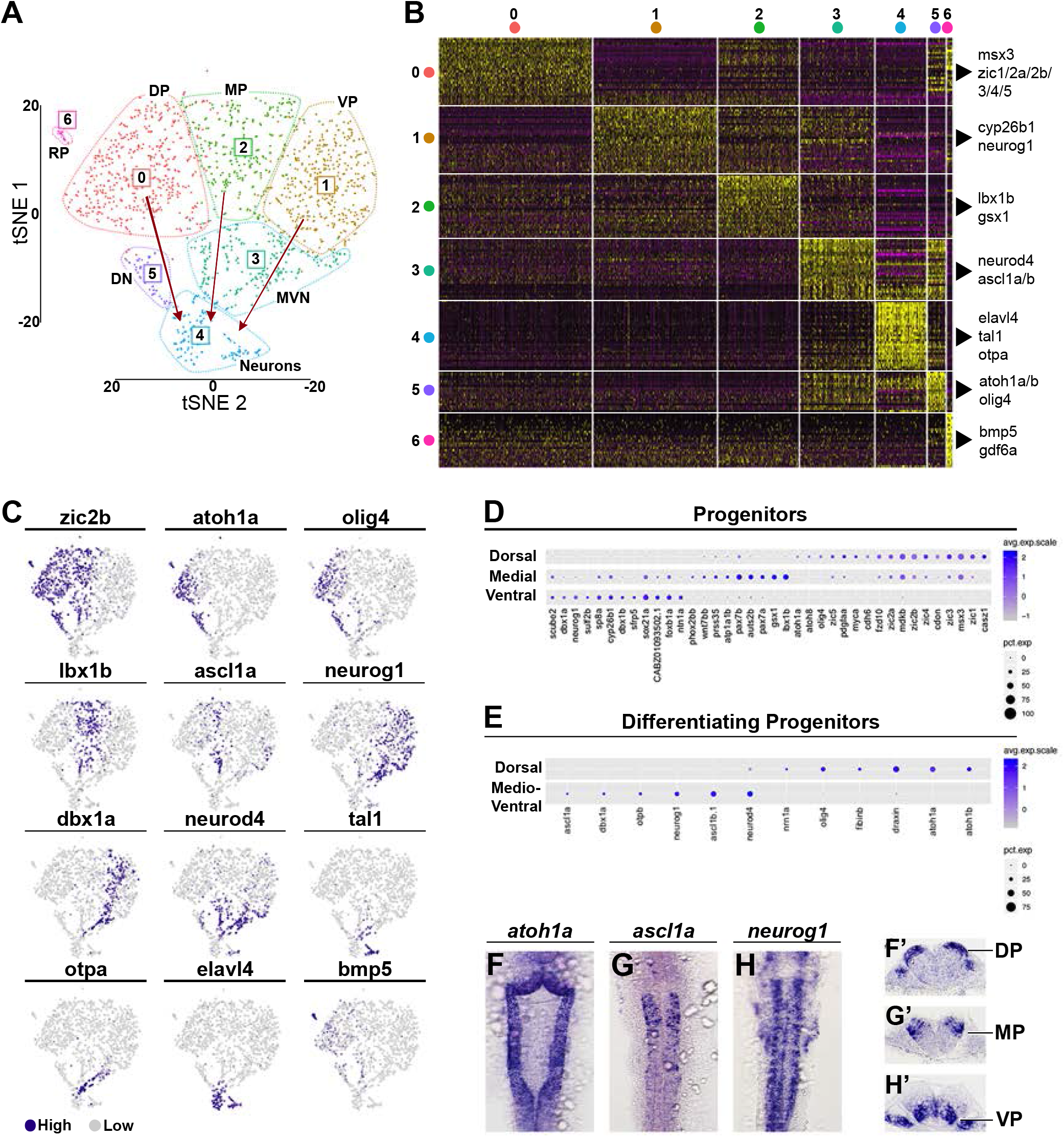
Cell population composition and signatures of the 24 hpf hindbrain. (A) Unsupervised tSNE plot subdivides hindbrain cells into 7 clusters. Dotted lines segregate each cluster, red arrowed lines indicate the direction of neurogenesis (Dp = Dorsal Progenitors, MP = Medial Progenitors, VP = Ventral Progenitors, DN = Dorsal Neurogenesis, MVN = Medio-Ventral Neurogenesis, RP = Roof Plate). (B) Heatmap of the top 30 genes significantly enriched in each cluster; representative gene names are shown close to each cluster. The full gene list is in Supplementary File 2. (C) tSNE plots showing the log normalised counts of selective representative genes. Colour intensity is proportional to the expression level of a given gene. (D) Dot Plot of genes with dorso-ventral restricted expression. (E) Dot Plot of factors with restricted expression in differentiating progenitors. Dot size corresponds to the percentage of cells expressing the feature in each cluster, while the colour represents the average expression level. Whole mount in situ hybridization showing the expression pattern of *atoh1a* (F, F’), *ascl1a* (G, G’) and *neurog1* (H, H’). (F’-H’) 40 μm hindbrain transverse section at the level of r4-r5/r5-r6.

**Figure 4.**
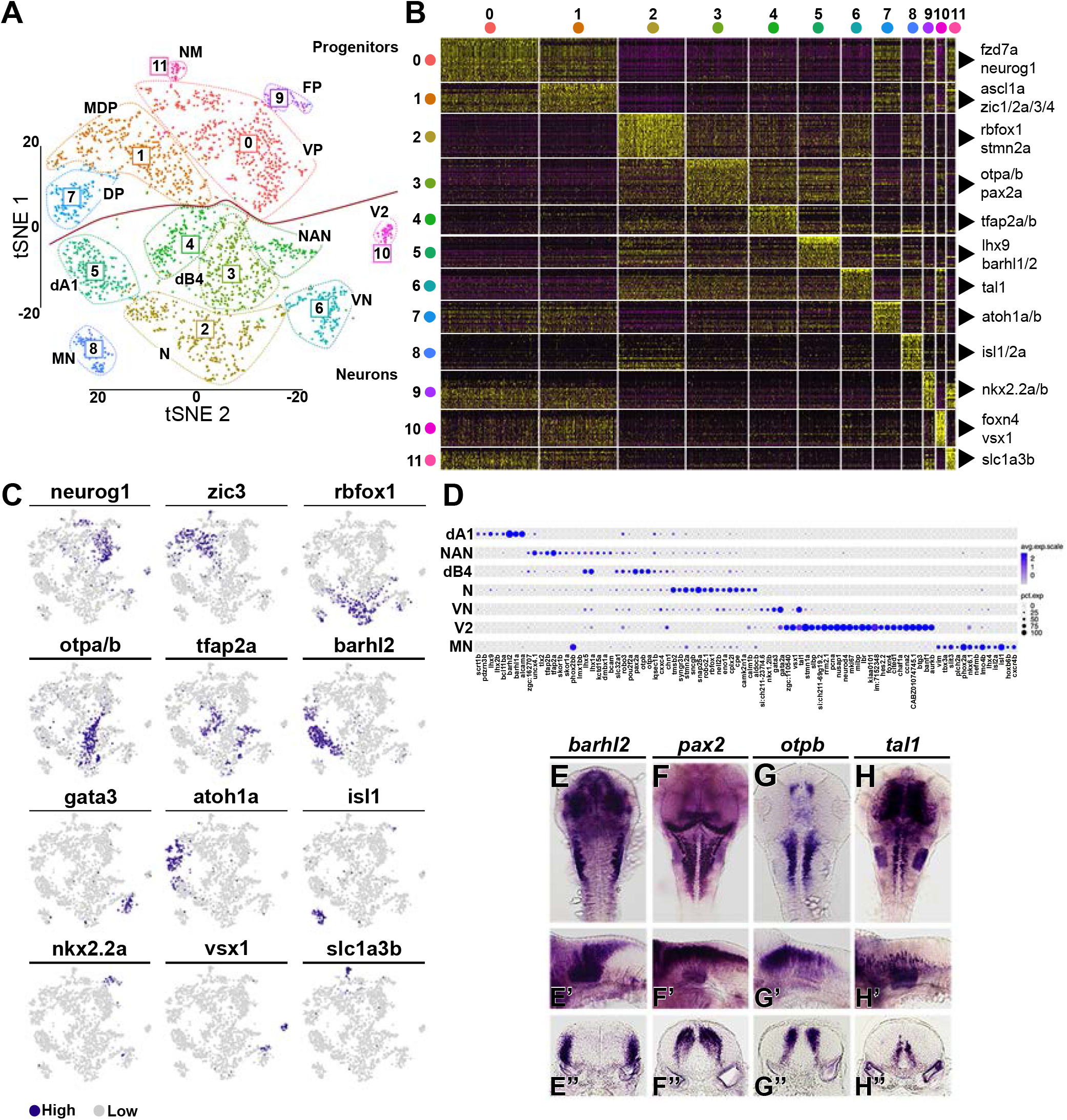
Neuronal complexity of the 44 hpf hindbrain. (A) Unsupervised tSNE plot subdivides cells into 12 clusters. Dotted lines segregate each cluster, red line divides progenitors from neurons (VP = Ventral Progenitors, MDP = Medio-dorsal progenitors, N = neurons, dB4 = GABAergic interneurons, NAN = NorAdrenergic Neurons, dA1 = dorsal neurons, VN = ventral neurons, DP = Dorsal Progenitors, MN = Motoneurons, FP = Floor Plate, V2 = lnterneurons, NM = Neuromast). (B) Heatmap of the top 30 genes significantly enriched in each cluster, representative gene names are shown close to each cluster. For the full gene list refer to Supplementary File 3. (C) tSNE plots showing the log normalised counts of selective representative genes. Colour intensity is proportional to the expression level of a given gene. (D) Dot Plot showing neuronal subtype molecular signature. Dot size corresponds to the percentage of cells expressing the feature in each cluster, while the colour represents the average expression level. Whole mount in situ hybridization showing the expression pattern of *barhl2* (E, E’), *pax2* (F, F’), *otpb* (G, G’) and *tal1* (H-H’). (E’-H’) 40 μm hindbrain transverse section at the level of r4-r5/r5-r6.

### Transcriptional signatures of hindbrain segments

The expression of known markers enables the identity of all clusters (C0-C8) at 16 hpf to be deduced (Fig.2A). At this stage, the main features that drive clustering of hindbrain cells using Seurat are segmental identity and dorsoventral (D-V) identity. We display the genes that distinguish the different clusters in a heatmap of the top 30 differentially-expressed genes (Fig.2B; Suppl. File 1) and show the expression level of selected genes in tSNE projection plots that relate them to the Seurat analysis (Fig.2C). Genes specifically expressed in different hindbrain rhombomeres (r), or in dorsal, medial or ventral domains, are listed in Fig.2D and Fig.2E, respectively.

tSNE projection plots with dorsal and ventral marker genes (Fig.2E) reveal the relationship between D-V identity and the clustering of cells in Seurat; see for example *zic2b* expression in Fig.2C which marks the dorsal part of the different hindbrain segments (Fig.2A). Cells from r2 and r4 co-cluster in C0-C1, where C0 cells are ventral and C1 cells are dorsal (Fig.2A). Seurat analysis did not discriminate r2 and r4 cells, suggesting strong transcriptional similarities, including *egfl6.1, fabp7a* and *sfrp5* expression. Cells from r3 are included in C0-C1, but form a discrete group that is marked, for example, by *egr2b* expression (Fig.2A, C). This clustering of r2, r3 and r4 cells reflects that genes including *hoxa2b, sfrp5* and *sp8a* are expressed in all three segments, whereas *egr2b, epha4a, sema3fb* and other markers are expressed in r3 cells (Fig.2C, D). Consistent with previous studies, r3 and r5 cells segregate to adjacent clusters, reflecting that they express some genes in common: in addition to the extensively-studied *egr2b* and *epha4a* genes, they express *timp2a, aldocb, smea3fb* and *myo1cb* (Fig.2D). r5 also shares transcriptional similarities with r6, which forms an adjacent cluster, including *mafba* (Fig.2C), *cryba2b, crygn2, lim2.1, col15a1b* and *gas6.* However, r7 cells (C4 ventral and C5 dorsal) do not cluster adjacent to r6 cells, reflecting that although some genes are expressed in both segments (for example, *hoxa3a, hoxb3a* and *tox3),* many other genes are expressed in one or the other, for example, *hoxd4a* (Fig.2C), *fabp7a, iratb, rbp5, rhbdl3* and *sp8a* in r7 (Fig.2D). r1 and midbrain-hindbrain boundary (MHB) cells which express known markers *(eng2a/b* (Fig.2C), *fgf8a, cnpy1* and pax2a) are found to cluster together in C6. Interestingly, floor plate cells which express *nkx2.2* (Fig.2C) are classified as part of cluster C6, suggesting some transcriptional similarities, but are clearly segregated from r1 /MHB cells and are adjacent to cells expressing ventral markers. Finally, roof plate cells (C8) form a discrete cluster adjacent to cells expressing dorsal markers. Since the organisation of clusters in Seurat favours local over global interactions, we also analysed the data using principal component analysis (PCA) which better represents global differences. We found that PCA clustered cells with distinct segmental identity in a similar way to Seurat, except that r6 rather than r5 was adjacent to the r2-r3-r4 cluster (data not shown).

As summarised in Suppl. Table 2 (Table 2.1), the transcriptome analyses have identified genes not previously described to have segmental expression in the hindbrain; these include *myo1cb* and *timp2b* in r3 and r5. In addition, we found genes for which expression data is available, but have not been tested functionally in the hindbrain; these include *sp8a* (strong in r4 and r7, weak in r2 and r3), *sfrp5* (r2-r4), and *wnt7aa* (r3-r7).

### Dorsoventral signatures of progenitors

Dorsoventral positional information is a key feature of the developing neural epithelium that underlies specification of neuronal cell types. Extensive molecular characterization has been carried out in the spinal cord (Delile et al., 2019; Gouti et al., 2015), but less widely for the hindbrain. At all stages analysed, progenitors were clustered based on their dorsoventral identity, reflecting that D-V patterning is established early and maintained during hindbrain neurogenesis. Seurat analysis at 16 hpf segregates cells into dorsal and ventral progenitors, as well as roof plate and floor plate (Fig.2A). tSNE projection plots with known markers (listed in Fig.2E) reveals that these are further subdivided into dorsal, medial and ventral domains. Seurat analysis at 24 hpf and 44 hpf clusters cells into discrete dorsal, medial and ventral populations, plus roof plate and floor plate (Fig.3A, Fig.4A). Selected genes that mark these different populations are presented in tSNE projection plots (Fig.3C, Fig.4C), and dot plots of relative expression levels (Fig.3D, Fig.4D).

The dorsal progenitors are identified by expression of known markers, which include *zic* genes (Elsen et al., 2008; Grinblat and Sive, 2001), *msx1b/3* (Miyake et al., 2012), *wnt8b* (Kelly et al., 1995), *olig3/4* (Tiso et al., 2009) and the proneural gene *atoh1* (Elsen et al., 2009). We find that dorsal progenitors also express *casz1, cdon, fzd10, myca, pdgfaa* and other genes (Fig.2B, E; Fig.3B-D; Suppl. Files 3, 4). In addition, *scrt1b, barhl1b* and *cdkn1cb* are dorsally restricted in 44 hpf progenitors (C7; Suppl. File 3). *atoh1a* (Fig.2E; Fig. 3F, F’; Fig.4C) and *atho8* (Fig. 3D) are expressed by both dorsal progenitors and differentiating cells, suggesting these are the main proneural genes contributing to dorsal neurogenesis. Additionally, the bHLH transcription factor *olig4* (also known as *zOlig3;* (Tiso et al., 2009)) is dorsally restricted, which has previously been shown to contribute to dorsal neural fate determination (Storm et al., 2009).

Medial progenitors share a few dorsally-(e.g. *zic* genes) and ventrally-expressed (e.g. *foxb1a)* factors, while uniquely expressing markers including *gsx1, pax7a/b* and *lbx1b* (Fig.2E; Fig. 3D; Suppl. File 1-2). *fabp7a,* a known astrocyte and radial glia marker, is enriched in the medial ventricular zone together with the newly identified factors *atp1b4* and *atp1a1b* (Fig.5D, Suppl. Fig.3K). This analysis further shows that the proneural genes *ascl1a* (Fig.3G, G’) and *ptf1a* (Suppl. Fig.3M) are expressed medially in the hindbrain, with expression overlapping with *neurod4* (Fig.3C).

**Figure 5.**
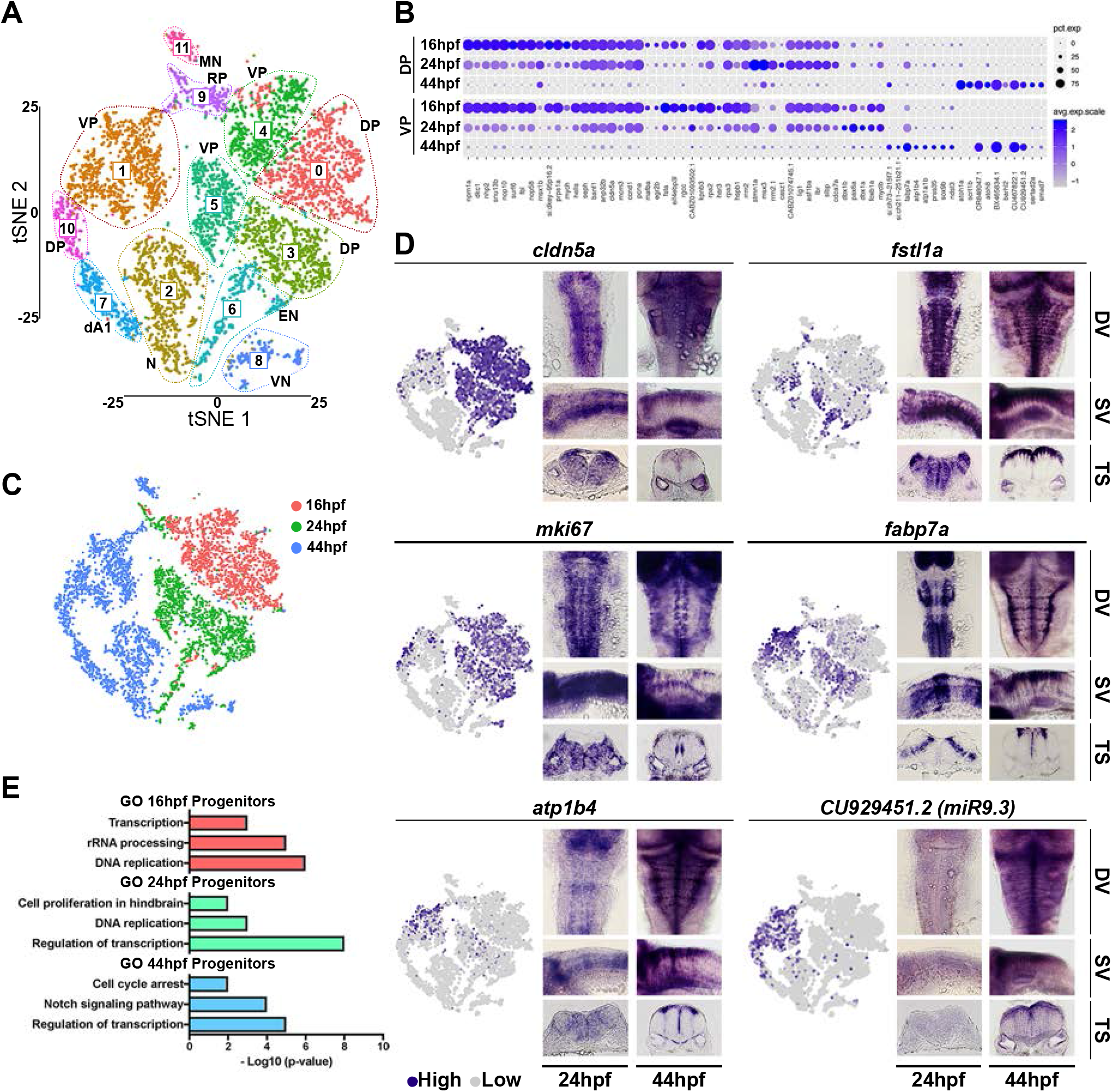
Analysis of aggregated 16 hpf, 24 hpf and 44 hpf data. (A) Unsupervised tSNE plot of cells from 16 hpf, 24 hpf and 44 hpf subdivides them into 12 clusters. DP = Dorsal Progenitors, VP = Ventral Progenitors, N = Neurons, EN = Early Neurogenesis 16-24hpf, dA1 = dorsal neurons, VN = ventral neurons, RP = Roof Plate, MN = Motoneurons. (B) Dot Plot showing molecular signature of dorsal and ventral progenitors at the three stages. Dot size corresponds to the percentage of cells expressing the feature in each cluster, while the colour represents the average expression level. The full gene list of top 30 significantly enriched factors is in Supplementary File 4. (C) tSNE plots with cells coloured based on their stage of origin: 16 hpf (pink), 24 hpf (green) and 44 hpf (blue). (D) tSNE plots showing the log normalised counts of representative genes. Colour intensity is proportional to the expression level of a given gene. Whole mount in situ hybridization showing the expression pattern of *cldn5a, fstl1a, mki67, fabp7a, atp1b4* and *CU929451.2 (miR9.3)* at 24 hpf and 44 hpf. Dorsal view (DV), side view (SV) and 40 μm hindbrain transverse section (TS) at the level of r4-r5/r5-r6 are shown for each gene. (E) Selected Gene Ontology (GO) terms at 16 hpf (pink), 24 hpf (green) and 44 hpf (blue) are shown. X-axis is −log10(p-value).

Lastly, ventral progenitors express the proneural gene *neurog1* (Fig.3H, H’) and *nkx6.2* (Fig.2E). Here, we found a ventral progenitor signature in which they express a unique set of transcription factors: *sox21a, foxb1a, sp8a* and *dbx1a/1b.* In addition, these cells express several signalling modulators: *sfrp5* (soluble inhibitor of Wnt signalling), *cyp26b1* (RA degradation), *scube2* (Shh long-range signalling), and *sulf2b* (heparan sulfate proteoglycans) (Fig.3), which may contribute to modulation of Wnt, RA and Shh levels that underlie neuronal cell type specification (Dessaud et al., 2008; Lara-Ramirez et al., 2013; Lupo et al., 2006; Ulloa and Marti, 2010).

### Characterization of neuronal complexity

Different neuronal subtypes are progressively generated from the dorsoventral progenitor domains. At 16 hpf, Seurat analysis identifies a single cluster (C7) expressing markers of neurogenesis (Fig.2A), and at 24 hpf and 44 hpf identifies distinct clusters that express early and late markers of neuronal differentiation (Fig.3A, Fig.4A). To determine whether the transcriptome of differentiating cells is similar or different at 16, 24 and 44 hpf, we aggregated the data and carried out Seurat analysis. Unsupervised clustering identifies 12 clusters and separates progenitors (C0, C1, C3, C4, C5, C10) and neurons (C2, C6, C7, C8, C11) (Fig. 5A). When cells are coloured by their developmental stage (Fig. 5C), we found that neurogenic cells at 16 hpf overlap with neurogenic cells at 24 hpf in cluster C6. Interestingly, they express the Activin-binding protein *fstl1a* and the Wnt antagonist *draxin* (Fig.5D), as well as transcriptional regulators including *ebf2* (Suppl. Fig.3J), *onecut1, scrt2* and *lin28a* (Suppl. File 4). In contrast, there is no overlap of neurogenesis at 16 hpf and 24 hpf with differentiating cells at 44 hpf (C2, C7, C8), consistent with the generation of new neuronal sub-types. There are also shifts in the transcriptome of progenitor cells which will be discussed below.

To characterise the neuronal complexity at 44 hpf, we classified neuronal sub-types based on (Hernandez-Miranda et al., 2017; Lu et al., 2015). Dorsal progenitors (atoh1a+, C7) generate dA1 excitatory interneurons (C5) in the hindbrain, a heterogeneous population that functions in sensory information processing (Hernandez-Miranda et al., 2017). *barhl1a, barhl2* (Fig.4C, E), *lhx2b* and *lhx9* are among their known markers, and in addition we find *alcama, bcl11ba* (BAF Chromatin Remodelling Complex), *pdzrn3b* and *scrt1b* (Fig.4D). Noradrenergic neuron (NAN) development is regulated by Phox2a and Phox2b (C4), and the proneural gene Ascl1a (C1) has been shown act upstream of Phox2 genes (Hirsch et al., 1998; Pattyn et al., 2000). In addition, Tfapa is important for activation of key NA enzymes (Holzschuh et al., 2003; Kim et al., 2001). These genes classify cluster C4, together with the transcription factors *dmbx1a, lhx1a/5, lmx1bb, tlx2* and *uncx4.1* which are co-expressed in these neurons (Fig.4D). Another class of hindbrain neurons are GABAergic inhibitory interneurons (dB4), here clustered in C3 (Fig.4A). For this class, *pax2, lhx1* and *lhx5* may constitute a transcription factor code (Burrill et al., 1997; Gross et al., 2002; Muller et al., 2002; Pillai et al., 2007). *slc32a1* (membrane protein involved in GABA uptake) is also highly expressed in this cluster. A subset of these cells coexpress *otpa/b* (Fig.4F-G), transcription factors involved in dopaminergic neuron specification (Fernandes et al., 2013). More ventrally, neurons are marked by *tal1* (Fig.4H) and *gata2a/3* expression (Fig.4C), resembling ventral neurons identified in the spinal cord (C6) (Andrzejczuk et al., 2018). A further cluster of ventral neurons is C10, which express *vsx1, tal1* and *foxn4,* defining this domain as V2 interneurons. Interestingly this cluster is characterized by expression of cell cycle factors (e.g. *ccna2, mki67, nusap1, pcna;* Suppl. Fig.4). vsx1-expressing cells in the hindbrain and spinal cord have been defined as non-apical progenitors, able to generate one excitatory (V2a) and one inhibitory (V2b) interneuron, and proposed to be a pool important for rapid generation of the sensory-locomotor circuit (McIntosh et al., 2017); their molecular signature is reported in Fig.4D. Motor neurons can be identified in C8 *(isl1, isl2, phox2a),* and in the hindbrain *lhx4, nkx6.1* and *tbx3a* are expressed in these cells (Fig.4D). A further neuronal cluster (C2) expresses a specific combination of genes (e.g. *aldocb, calm1b, camk2n1a, rbfox1;* Fig.4C, D), but could not be classified. C11 consists of neuromast cells that were present in the dissected tissue and had not been removed bioinformatically. Our transcriptome atlas thus gives new insights into factors expressed in different hindbrain neuronal subtypes (Fig.4D; Suppl. File 3).

### Transcriptional shift of hindbrain progenitors

In addition to temporal differences in expression of neurogenic markers, Seurat analysis found changes in the transcriptome of progenitor cells (Fig.5A, C). 16 hpf and 24 hpf progenitors are in distinct clusters, but in close proximity in tSNE space, whereas 44 hpf progenitor cells are further apart (Fig. 5C). Analysis of the top 30 significantly enriched genes highlights transcriptional similarities and differences between progenitors (Suppl. File 4). Both dorsal and ventral progenitors have a similar transcriptional signature at 16 and 24 hpf. The genes enriched in these cells are reported in Fig.5B; among them *fsta* (Fig.6J), *cldn5a* (Fig.5D), *pax6a* and proliferative markers (Fig.5D) are strongly expressed.

**Figure 6.**
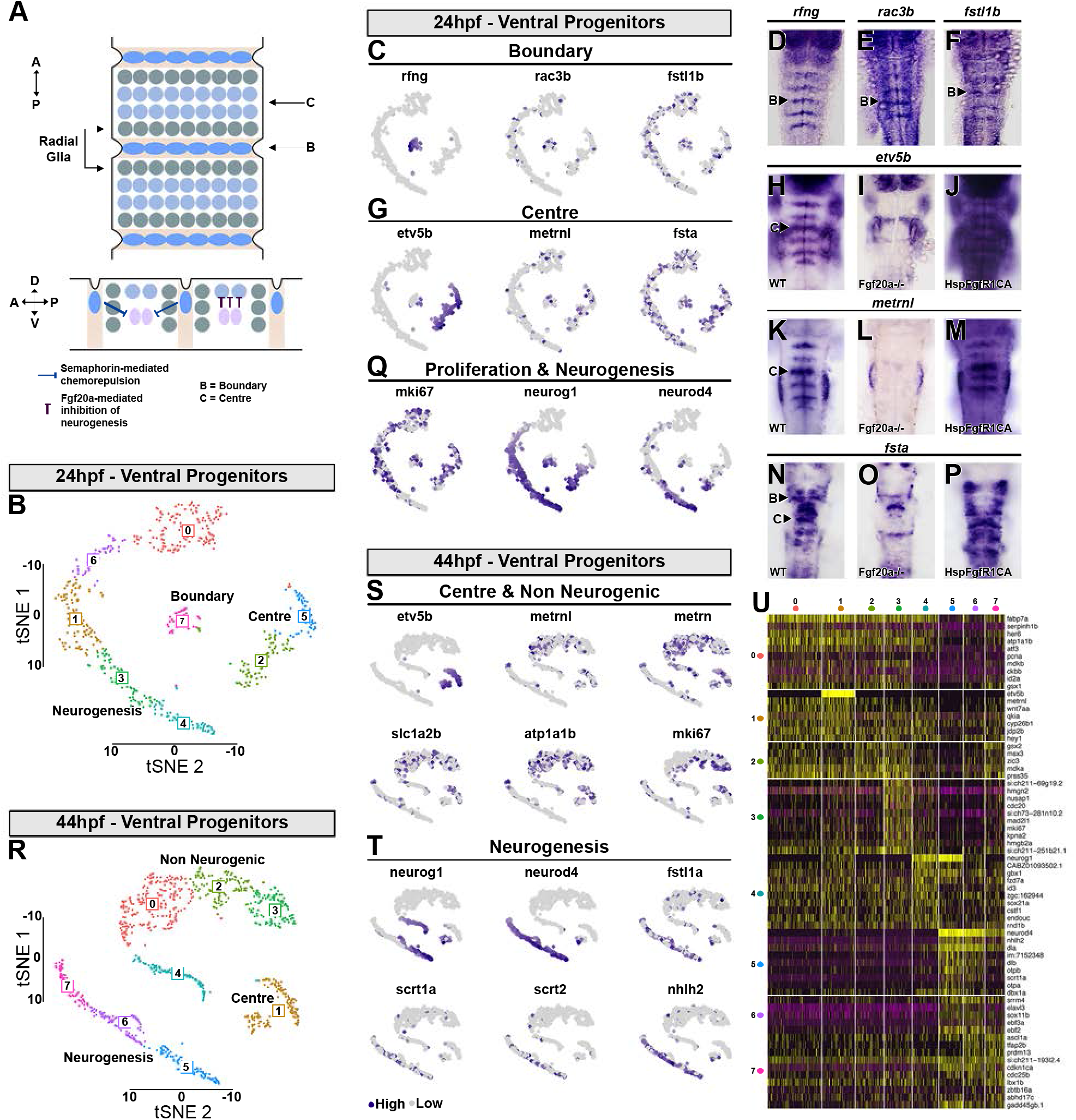
Transcriptional signature of boundary cells and segment centre progenitors. (A) Schematic drawing representing anterior-posterior organization within hindbrain segments. Boundary cells are in cyan, neurogenic progenitors in grey and segment centre cells in light blue. Below is a side view showing the role of boundary cells in maintaining fgf20a neurons (pink) at the centre of each segment, mediated by semaphorins. Fgf20 signaling maintains undifferentiated progenitors. (B) Supervised clustering of 24 hpf ventral progenitors. 8 clusters are identified: C0, C6, C1 are progenitors; C3, C4 are neurogenic domains; C7 are boundary cells; and CZ, CS are segment centre progenitors. tSNE plot showing the expression distribution of boundary (C), segment centre (G) and proliferation and neurogenic genes (Q). Whole mount in situ hybridization of boundary (D-F) and segment centre genes (H, K, N). Segment centre-specific gene expression is dependent on Fgf20 signalling, as fgf20a-/- embryos have loss of *etv5b* (I), *metrnl* (L) and *fsta* (O) expression, whereas constitutive activation of FgfR1 induces their ectopic expression (J, M, P). (R) Supervised clustering of 44 hpf ventral progenitors. 8 clusters are identified: C3, C2, C0 are progenitors; C4, C7, C6, C5 are neurogenic domains; C1 cells are segment centre progenitors. tSNE plot showing the expression distribution of segment centre and non-neurogenic genes (S) and neurogenic genes (T). (U) Heatmap of the top 10 genes enriched in each cluster.

Gene ontology terms associated with the top 30 genes enriched in C0-C4 (16 hpf) and C3-C5 (24 hpf) highlight the proliferative property of these progenitors (Fig.5E). A drastic reduction in proliferation has taken place by 44 hpf. For example, *mki67, nusap1, ccnd1* and *cdca8* are widely expressed in the early hindbrain, whereas they are restricted to a small proportion of dorsal progenitors and vsx1-expressing cells at 44 hpf (Suppl. Fig.4; Fig.5D). In addition, genes associated with cell cycle arrest *(cdkn1ca, cdkn1cb)* and Notch signalling are increased at 44 hpf (Fig.5E). Glial cells become apparent at 44 hpf in the medio-ventral progenitor pool marked by *fabp7a,* and we find they also express *atp1b4* and *atplalb* (Fig.5D, Suppl. Fig.3K). Furthermore, *miR9* loci are detected only at 44 hpf *(miR9.1 CR848047.1, miR9.3 CU929451.2, miR9.6 CU467822.1)* (Fig.5B, D), when they play a key role in the timing of neurogenesis (Coolen et al., 2013; Coolen et al., 2012). Overall, this analysis highlights that there are significant changes in gene expression in progenitors between 24 hpf and 44 hpf in the developing hindbrain.

### Boundary cell and segment centre progenitors

During hindbrain development in zebrafish, proneural gene expression becomes confined to zones flanking the segment boundaries, with low expression in boundary cells and segment centres (Amoyel et al., 2005; Cheng et al., 2004; Gonzalez-Quevedo et al., 2010). The formation of these non-neurogenic zones (Fig.6A) involves Notch (Cheng et al., 2004) and Yap/Taz (Voltes et al., 2019) at boundaries, and Fgf20 signaling at segment centres (Gonzalez-Quevedo et al., 2010). These distinct progenitor populations were not identified by unsupervised clustering, because this is dominated by the large differences in the transcriptome during dorsoventral patterning and differentiation. We therefore used supervised clustering with known markers to reveal the transcriptional signature of the neurogenic and non-neurogenic cell populations.

We bioinformatically isolated 24 hpf ventral progenitors and used *rfng* (boundary), *etv5b* (segment centre), and *neurog1* and *neurod4* (neuronal differentiation) to drive clustering. Seven sub-clusters were obtained (Fig.6B), with C0, C6, C1, C3, C4 forming a continuum and presenting a neurogenic gradient (low to high along tSNE1; Fig.6Q). We found that boundary cells that express *rfng* (Fig.6D, C7) express some previously known markers: *rasgef1ba* (Letelier et al., 2018), the Rho GTPase *rac3b* which maintain sharp borders (Fig.6E; Letelier et al., 2018) and the RA-degrading enzyme *cyp26b1* which may contribute to regulation of neurogenesis (Gonzalez-Quevedo et al., 2010). In addition, we find new genes with expression enriched at boundaries including *rnd2, dystroglycan 1* (Parsons et al., 2002) and the BMP inhibitor *follistatin 1b* (Fig.2F; Dal-Pra et al., 2006). There is also higher expression of *cyclin A2* (Suppl. File 5) and *cyclin D1* (data not shown) which could regulate distinct proliferative properties. Finally, the transcription factors *pax6a* (Kleinjan et al., 2008) and *nr2f2* (Love and Prince, 2012) have wide hindbrain expression with some boundary enrichment. Thus, we identified a distinct set of factors present in boundary cells.

At each segment centre, cells upregulate the Fgf-direct target *etv5b* (Esain et al., 2010; Gonzalez-Quevedo et al., 2010) which we used to drive clustering of 24 hpf progenitors. etv5b-expressing cells are in two clusters, C5 and C2. In C2 there is transcriptional overlap of *etv5b* with *neurog1, ascl1b.1* and *neurod4* (Fig.6G, Q) while in C5 all cells are non-neurogenic. The overlap in C2 reflects that at 24 hpf, *etv5b* is expressed in stripes located at the centre of each segment (Fig.6H) but neurogenic gene expression has yet to be down-regulated (Fig.3H; Gonzalez-Quevedo et al., 2010). Many of the genes in C5 have an unknown expression pattern (Suppl. Table 2), but previous work (Gonzalez-Quevedo et al., 2010) showed that segment centre marker genes are under Fgf20 control. We therefore performed a bulk RNA-seq experiment comparing wild-type to *fgf20a* mutant hindbrain. *metrnl* and *fsta,* both present in C5 (Fig.6G, K-N), were among the downregulated genes (Suppl. Fig.5B; Suppl. Table 3). However, *etv5b* was not found, which likely reflects that it is expressed both in the hindbrain and adjacent otic vesicle and cranial ganglia; Suppl. Fig.5A). We therefore also profiled transgenic hindbrains expressing heat-shock induced constitutively active FgfR1 (Tg(hsp70:ca-fgfr1)) and compared to heat-shocked controls. This screen found *etv5b, metrnl* and *fsta* among the top genes induced by Fgf signalling (Suppl. Fig.6; Suppl. Table 4). In situ hybridization confirmed that Fgf20 signalling is both necessary for expression of *etv5b, metrnl* and *fsta* in segment centres (Fig.6H-P). *metrnl* encodes a cytokine with an unknown receptor. Since the related meteorin gene (metrn) is implicated in gliogenesis in other contexts (Lee et al., 2010; Nishino et al., 2004), it is a candidate to promote glial cell differentiation that occurs at segment centres. Interestingly, *fsta* is also expressed by boundary cells, and thus correlates with non-neurogenic progenitors. Overall, we found a limited number of genes exclusively expressed by boundary or centre progenitors, while the majority of transcripts are expressed in a scattered pattern across the two cell populations (Suppl. File 5).

At 44 hpf, neurogenic zones are fully refined but *rfng* is no longer detected. We therefore only used *etv5b* and *neurog1+neurod4* to drive clustering. At this stage, the number of neurogenic etv5b-expressing cells has greatly decreased (Fig.6S-U) and they are expressing many ventral progenitor genes. *metrn* and *metrnl* are expressed in a similar pattern to *slc1a2b, atp1a1b* and other glial markers, further suggesting that the Metrn family could play a role in hindbrain gliogenesis. neurod4-expressing cells are separated from the remaining progenitors (Fig.6R, T) and present a unique signature (Fig.6U). Interestingly *fstl1a* (Fig.5D), the transcription factors *scrt1a, scrt2* (Fig.7C, E) and *nhlh2* are co-expressed in these cells (Fig.6T).

**Figure 7.**
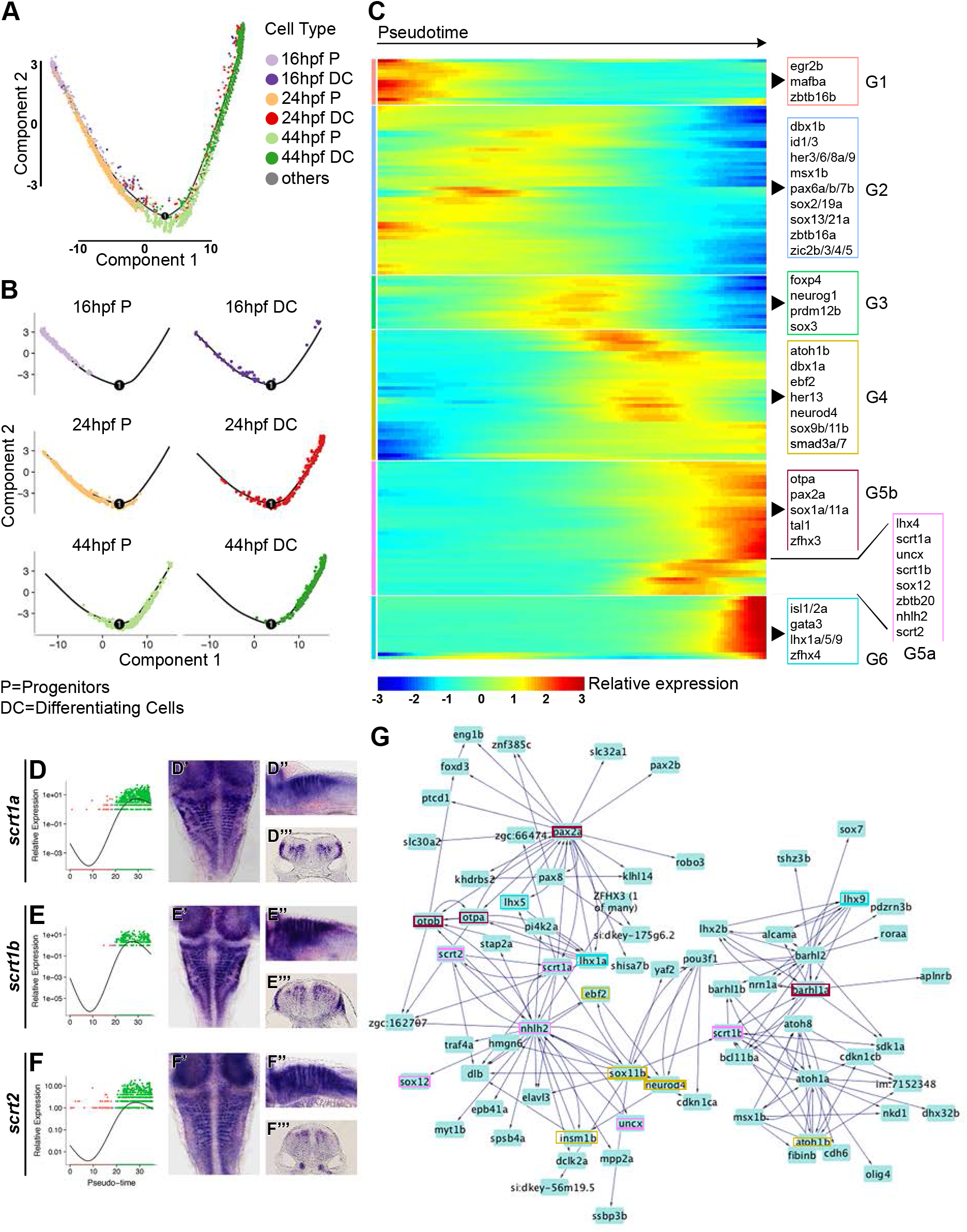
Analysis of transcription factor expression during hindbrain neurogenesis. (A) Pseudotemporal ordering of 16 hpf, 24 hpf and 44 hpf hindbrain progenitors (P) and differentiating cells (DC) by Monocle 2. Cells are coloured by developmental stage and differentiation state. (B) Individual pseudotemporal plots representing progenitors and differentiating cells distribution at each developmental stage. (C) Heatmap showing TFs clustered by pseudotemporal expression pattern (q values<0.01). Pseudotime ordering goes from left (progenitor state) to right (differentiated neurons). Selected transcription factors are shown for each group (G1-G6). The full gene list is in Supplementary File 7. (D-F) Expression of *scrt1a, scrt1b* and *scrt2* during pseudotime. Whole mount in situ hybridization at 44 hpf for *scratch* genes is shown in dorsal view (D’-F’), side view (D”-F”) and hindbrain sections (D”’-F”’). (G) Using GENIE3, a directed network of interactions was predicted among the genes in the 44 hpf scRNA-seq data set. The *scratch* genes network was view and extracted in Cytoscape; boxes highlight TFs present in the above heatmap, and colours match the group of origin in (C).

### Transcription factors temporally regulating hindbrain neurogenesis

To illustrate developmental insights that can be extracted from the single cell RNA-Seq data, we focused on transcription factors (TFs) (AnimalTFDB3.0 database; Zhang et al., 2012) and inferred their potential contribution to hindbrain neurogenesis. We used the aggregated data set (Fig.5A) and performed pseudotime analysis using Monocle 2 (Qiu et al., 2017; Trapnell et al., 2014b), which orders cells uniquely on the similarity of their global TF expression profiles. This created a pseudotime trajectory with three discrete cell states (Suppl. File 6). The root of the trajectory was defined as the state containing the majority of the 16 hpf progenitor cells. The three states are characterized by the expression of: *sox2, egr2b, mafba, zic* genes, *pax6a/b* and *zbtb16a/b* among others for the progenitor state; *sox3, neurog1, atoh1a, dbx1a, gsx1, lbx1b* in the intermediate differentiation state; *atoh1b, neurod4, isl 1, vsx1, tal1, pax2a* and other neuronal TFs have high expression in the final state (Suppl. File 6). Along the trajectory, cells are ordered largely based on developmental stage of origin and state of differentiation (Fig. 7A, B). 16 hpf and 24 hpf progenitors are mainly at the start of the trajectory, followed by 44 hpf progenitors. 16 hpf differentiating cells present a TF expression pattern that mostly resembles 24 hpf progenitors, with the exception of few cells found at the end of the trajectory, while 24 hpf and 44 hpf differentiating cells largely overlap (Fig.7B). These data further suggest transcriptional changes in early versus late hindbrain progenitors.

To identify a temporal cascade of TFs potentially involved in neurogenic cell-fate decisions, we mapped TFs that significantly varied in their pseudo-temporal expression pattern, and clustered them according to their expression dynamic (Fig. 7C; Suppl. File 7). This analysis highlights multiple discrete shifts in TF expression occurring during hindbrain neurogenesis. Six distinct patterns were identified, where the first has high expression at the beginning of pseudotime, and the others progressively shift until reaching a peak of expression of neuronal markers. The first group (G1) includes *egr2b* and *mafba,* which are involved in segmental identity of progenitors and rapidly down-regulated at the onset of differentiation. In the next group (G2) are genes expressed in progenitors but not down-regulated until later in pseudotime, and implicated in the maintenance of the progenitor fate and/or inhibition of neurogenesis. Among them, *zic* and her genes promote neural progenitor identity and inhibit differentation (Bae et al., 2005; Coolen et al., 2012; Nyholm et al., 2007; Scholpp et al., 2009), *id* genes encode negative regulators of proneural bHLH proteins (Ellis et al., 1990; Garrell and Modolell, 1990; Ling et al., 2014) and *zbtb16a (plzfa)* inhibits neurogenesis (Sobieszczuk et al., 2010). The following group of genes (G3) with shifting expression in pseudotime are: *sox3* which has initial constant expression followed by a drop in differentiated cells; *neurog1* (reviewed by Bertrand et al., 2002); *prdm12b,* a regulator of V1 interneuron fate decision (Thélie et al., 2015; Zannino et al., 2014); and *foxp4* that is promotes detachment of differentiating cells from the neuroepithelium (Rousso et al., 2012). *atoh1b* and *neurod4* are in the next step of the cascade (G4) together with *ebf2,* a factor that acts downstream of proneural genes and necessary for initiation of migration and neuronal differentiation (Garcia-Dominguez et al., 2003). In the next group, a subset of genes initiates expression that then declines late in pseudotime (G5a). They include *zbtb20* that functions during corticogenesis as in the generation of layer-specific neuronal subtypes (Tonchev et al., 2016), and the less-studied *uncx, nhlh2, lhx4* and *sox12.* Furthermore, members of the *scratch* family *(scrt1a/b/2)* has a similar dynamic pattern, with enrichment within the neurogenic zone and some dorso-ventral differences: *scrt1a* and *scrt1b* are expressed ventral and dorsal (Fig. 7D-E) while *scrt2* is only found ventral (Fig. 7F). These genes have been implicated in the onset of neuronal migration (Itoh et al., 2013; Paul et al., 2014). Also classified in this group, but with a later onset of expression that does not decline (G5b) are transcription factors implicated in neuronal specification *(otpa, tal1, pax2a).* The final group of genes with an onset of expression late in pseudotime (G6) also encode regulators of neuronal identity *(isl1/2a, gata3, lhx1a/5/9)*.

To further explore TFs role in hindbrain neurogenesis we used a complementary approach that does not relay on pseudotemporal ordering. A gene regulatory network (GRN) was created using GENIE3 (Huynh-Thu et al., 2010), which uses a Random Forest machine-learning algorithm to predict the strength of putative regulatory links between a target gene and the expression pattern of input genes (transcription factors). A GRN was produced for each individual stage (data not shown), and here we focus on 44 hpf since it is relevant for late steps of neurogenesis. To focus on the predictions with higher significance, we applied a threshold of >0.025 of important measure (IM) and these interactions were analysed in Cytoscape (Shannon et al., 2003) (Suppl. Table 5). This cut-off recovered 4637 total interactions that constitute a valuable resource to guide future *in vivo* functional validations. Given the complexity of the network we extracted a submodule to exemplify its predictive potential. We interrogated the network to specifically analyse *scrt* genes during neurogenesis, and extracted their closest neighbours (Fig.7G). This network module predicts interconnections between genes in G5a, G5a and G4. *scrt1a* and *scrt2* are found in a feedback loop with *nhlh2,* and upstream of neurogenic factors *(neurod4, elavl3, otpa/b,* and *pax2a),* while *scrt1b* is connected to *atoh1a/b, atoh8,* and *barhl1a/b*.

## DISCUSSION

The single cell transcriptome atlas that we present here is a resource for further investigation of mechanisms that regulate neurogenesis and other aspects of hindbrain development. We analysed the transcriptome of hindbrain cells prior to (16 hpf), during (24 hpf) and after (44 hpf) the patterning of neurogenesis to form discrete neurogenic and non-neurogenic zones within segments. We used unbiased methods to cluster cells based on transcriptional differences, and identified genes that mark distinct hindbrain segments, cell types along the dorsoventral axis, and neuronal differentiation. By comparing our findings with previous studies, we have created an annotated list of genes that indicates which are previously known and which are novel markers, as also highlighted in the relevant Results section.

Seurat analysis at 16 hpf clustered cells based on segment-specific gene expression and gave a global picture of differences in the transcriptome of distinct segments. The organisation of clusters from r2 to r6 suggests that neighbouring segments have a similar transcriptome, but with a significant difference between odd- and even-numbered segments. This is consistent with previous studies showing nested expression of *hox* genes that regulate anterior-posterior identity (reviewed by Alexander et al., 2009; Tümpel et al., 2009), and the role of *egr2* in regulating gene expression in r3 and r5 that confers distinct properties from r2, r4 and r6 (Voiculescu et al., 2001). In contrast, r7 cells do not cluster adjacent to r6 cells, suggestive of a distinct identity which may reflect that it is a transitional zone to the anterior spinal cord.

We find major differences in gene expression in differentiating neurons at 16 hpf and 24 hpf compared with 44 hpf, as expected from the generation of distinct neuronal cell types at different times. Our analyses reveal new genes that are co-expressed with known markers of neuronal cell types that form along the dorsoventral axis. In addition to transcription factors, these include modulators of the Shh, RA and Wnt pathways. Interestingly, many differentiating neurons at all stages express *fstl1a,* suggesting a potential role of BMP inhibition. The generation of different neuronal cell types at 44 hpf compared with 16 hpf and 24 hpf is accompanied by changes in gene expression in progenitor cells at these stages, including proliferation markers and miR9 microRNAs. By carrying out pseudotime analysis, we inferred progressive changes in gene expression during the differentiation of progenitor cells to neurons. These data suggest a cascade in which genes that define segmental identity are rapidly down-regulated, followed by factors that maintain progenitor cells, in turn followed by upregulation of genes required for neuronal migration and transcription factors that define neuronal identity. We also analysed transcription factor expression using an algorithm to predict gene regulatory networks. We focussed on *scrt* family genes that regulate neuronal migration, and this found potential relationships with proneural factors and regulators of neuronal identity. We envisage that investigators can interrogate the network for other TFs of interest to guide biological hypotheses and phenotypic screening of specific mutants.

One motivation for this study was to find genes that mark the distinct neurogenic and non-neurogenic zones that are established in the zebrafish hindbrain. These features are not found in the unbiased analysis, as this is dominated by the greatest transcriptomic differences. We therefore used known markers of hindbrain boundary cells, neurogenic cells and segment centres to drive clustering of the progenitor population. In addition, we carried out RNA-seq analyses after manipulation of Fgf pathway activation which inhibits neurogenesis at segment centres. These analyses identified novel signaling factors, most notably follistatin and meteorin family members expressed in boundary cells and/or segment centres that are candidates to inhibit neurogenesis or promote gliogenesis. The single cell transcriptome data will enable investigators to extract information on other specific cell populations by this approach.

## MATERIALS AND METHODS

### Maintenance of zebrafish strains and husbandry

Zebrafish embryos were raised at 28.5° C or 25° C depending on the required stage (Westerfield, 2007). Embryos were staged according to hour post fertilization (hpf) and morphological criteria (Kimmel et al., 1995). The zebrafish work was carried out under a UK Home Office Licence under the Animals (Scientific Procedures) Act 1986 and underwent full ethical review.

### Mutant strains and heat shock treatment

*fgf20a* (dob) mutant embryos (Whitehead et al., 2005) were obtained from homozygous mutant in-crosses. Transgenic *Tg(hsp70:ca-fgfr1)* embryos are heterozygotes from outcrosses (Gonzalez-Quevedo et al., 2010; Marques et al., 2008). To induce constitutively active Fgfr1, *Tg(hsp70:ca-fgfr1)* embryos at 22 hpf were heat shocked for 30 min at 38.5° C and then incubated for 2 h at 28.5° C. Since around 50% of the embryos are carrying the transgene, controls and treated embryos were collected from the same heat-shocked clutch, avoiding any issue with differences in genomic background and changes in gene expression due to the heat shock treatment. After mRNA extraction, qPCR was performed to identify properly dissected tissues and discriminate between controls and *fgfr1* over-expressing tissues.

### Whole-mount in situ hybridization

For whole-mount in situ hybridization, embryos or explants were fixed in 4% PFA overnight at 4°C, or 4 h at room temperature, and kept in methanol at −20°C prior to processing. Some probes have been previously described: *neurog1* and *neurod4* (Alexander et al., 2009; Gonzalez-Quevedo et al., 2010), *pax2* (Krauss et al., 1991), *rfng* (Cheng et al., 2004), *etv5b* (cb805, ZFIN), *metrnl* (MPMGp609H2240Q8, RZPD), *sox3* (EST clone: IMAGp998H108974Q). Additional probes were generated from cDNA of 20-44 hpf embryos. A forward primer was used together with a reverse primer with a T7 promoter site (5’gaaatTAATACGACTCACTATAGg3’) for amplification; see Table 1. Digoxigenin-UTP labelled riboprobes were synthesised and *in situ* hybridization performed as previously described (Xu et al., 1994). After BCIP/NBT colour development, embryos were re-fixed for 30 min, cleared in 70% glycerol/PBS, and mounted to view the dorsal or lateral side. For transverse sections, embryos were extensively washed in PBST prior to mounting in 4% agarose/water. Embryos were sectioned using a Vibratome (Lecia VT1000 S), generating transverse sections of a thickness of 40 μm. Imaging was carried out with a Zeiss Axioplan2 with an Axiocam HRc camera.

**Table 1.**
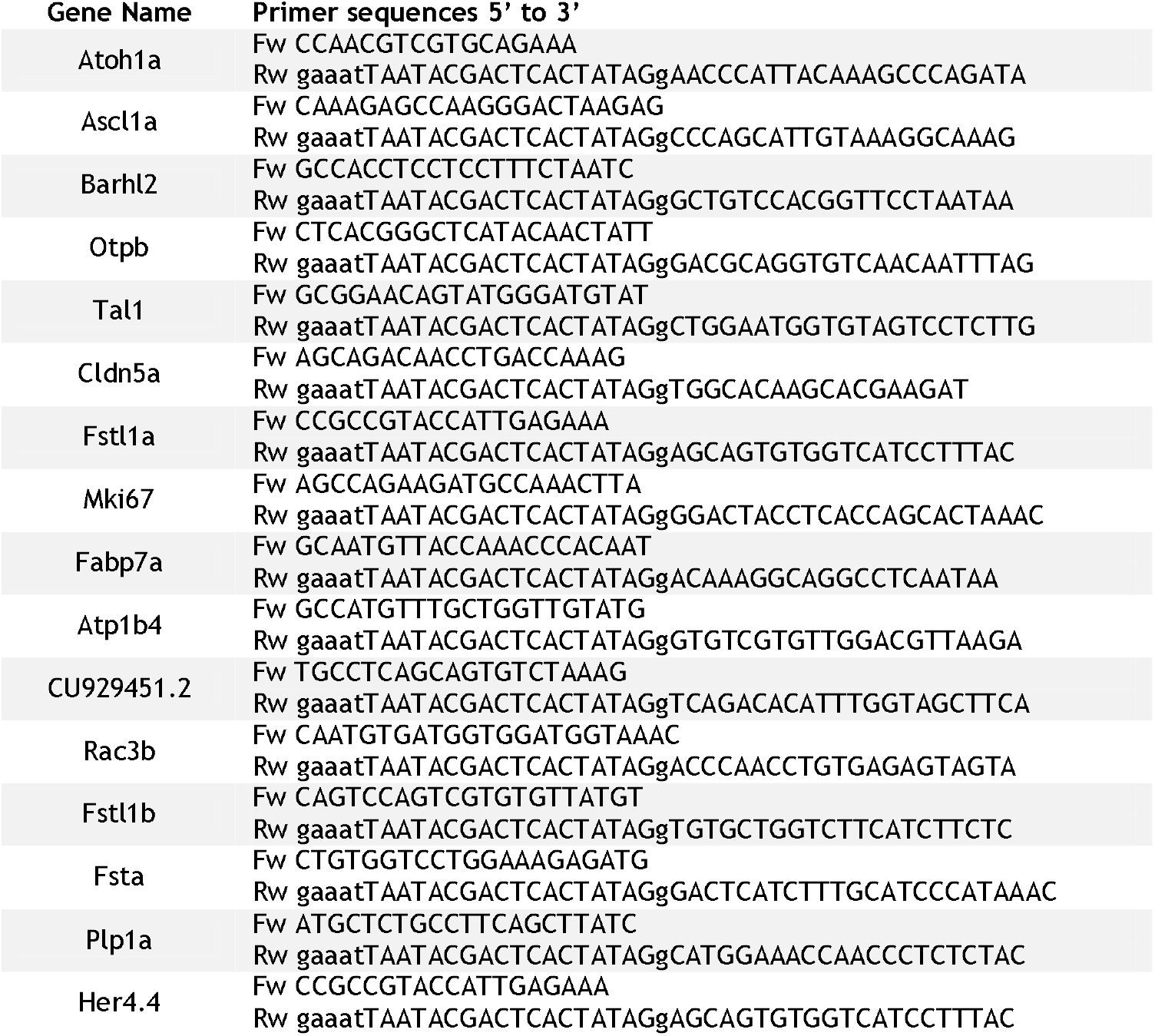
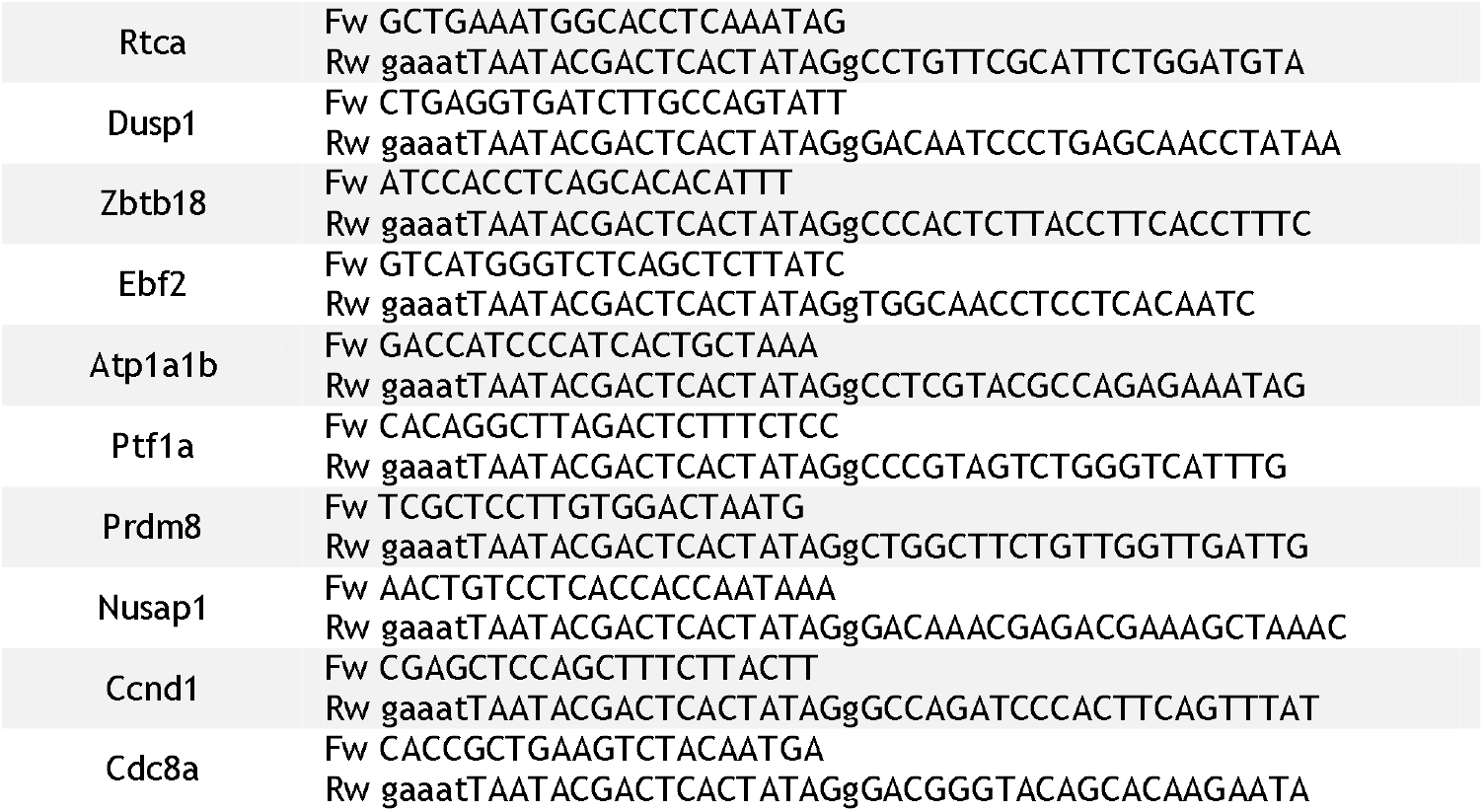
Primer sequences for antisense probe generation.

### Hindbrain dissection

Embryos at the desired stage were decorionated and de-yolked in DMEM with high Glucose, no Glutamine, no Calcium (11530556, Gibco); hindbrains were micro-dissected using 0.33 mm micro-fine sterile needles. Dissected tissues were kept in DMEM until further processed. For RNA-seq a single hindbrain tissue was collected in an individual tube and the quality of the dissection evaluated by qPCR (data not shown). For scRNA-seq, around 40 tissues per stage were pooled and immediately processed for cell dissociation.

### RNA extraction, cDNA preparation and qPCR

RNA was isolated using Quick-RNA Microprep kit (Zymo Research) and eluted in 15 μl (Lan et al., 2009). To evaluate the quality of dissection 3 μl of RNA was reverse transcribed using SuperScript™ III Reverse Transcriptase (ThermoFisher Scientific), and the remainder stored at −80° C until processed. Primers for target genes were designed with PrimerQuest (IDT). qPCR was performed using QuantStudio 3 (ThermoFisher Scientific) with SYBR green Platinum™ SYBR™ Green qPCR SuperMix-UDG (ThermoFisher Scientific) master mix. The ΔΔCt method was used to calculate gene expression (Livak and Schmittgen, 2001). β-actin was used as reference gene. Primers used are listed in Table 2. Samples without contamination were processed for RNA-seq.

**Table 2.**
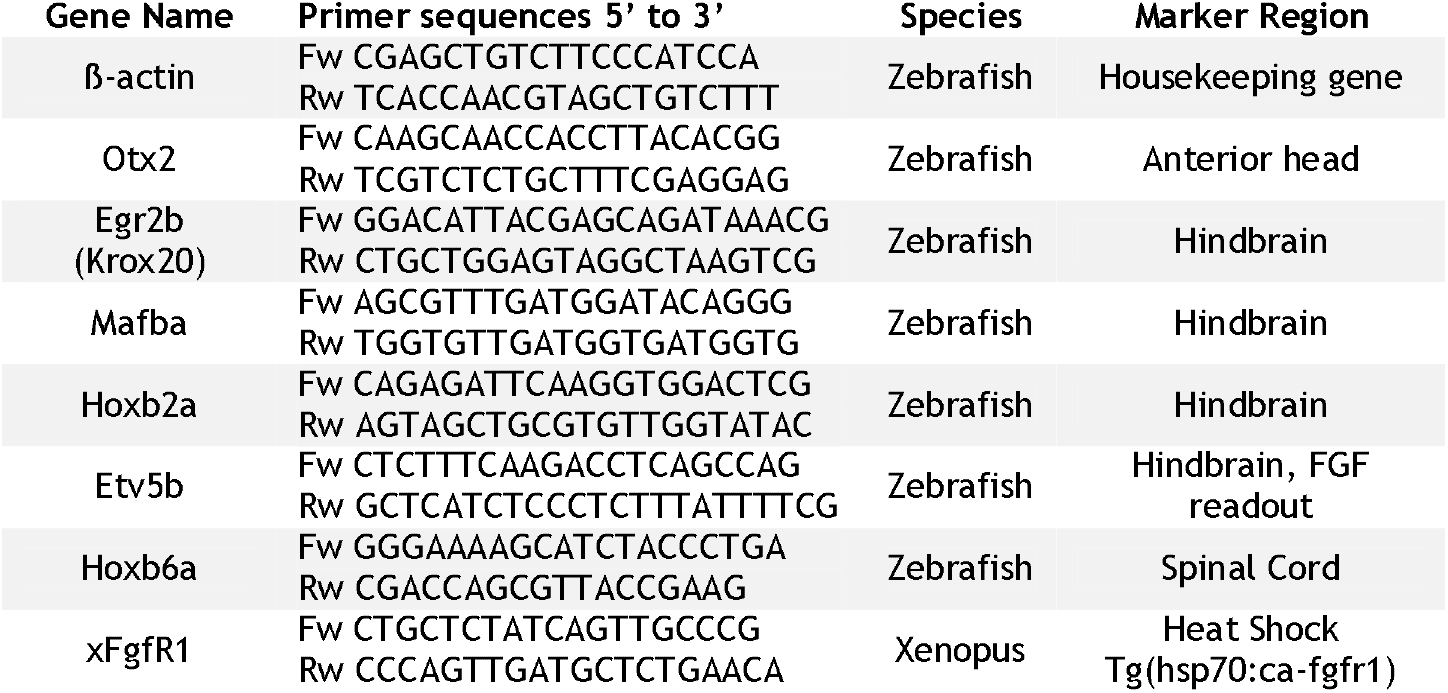
qPCR primers.

### Library Preparation and RNA-sequencing

Libraries for the *fgf20a-/-* experiment were prepared with the Ovation® RNA-Seq System V2 (7102, NuGEN) for cDNA amplification, followed by NexteraXT (Illumina) for library preparation. These libraries were sequenced on the HiSeq 2000 (Illumina), with paired-end 75 bp reads. Libraries for the constitutive active Fgfr1 experiment were prepared with the Clontech SMARTer kit (634926, TaKaRa) for cDNA amplification, followed by NexteraXT (Illumina) for library preparation. These libraries were sequenced on the HiSeq 4000 (Illumina), with single ended 75 bp reads.

### Sequence alignment and analysis of differentially expressed genes

The quality of the samples was assessed using FastQC. Reads were aligned against zebrafish genome GRCz10 and Ensembl release 86 transcript annotations using STAR v2.5.1b (Dobin et al., 2013) via the transcript quantification software RSEM v1.2.31 (Li and Dewey, 2011). Gene-level counts were rounded to integers and subsequently used for differential expression analysis with DESeq2 v1.20.0 (Anders and Huber, 2010) using default settings. Differential expression results were thresholded for significance based on an FDR<=0.01, a fold-change of +/− 2 and a minimum normalized count of >30 in all contributing samples from at least one of the replicate groups being compared. Heatmaps were created using rlog transformed count data, scaled across samples using a z-score.

### Preparation of single cells from zebrafish hindbrain

Around 40 hindbrain tissues per stage (16 hpf, 24 hpf, 44 hpf) were dissected as described above. The samples were incubated with FACS max cell dissociation solution (T200100, Amsbio) supplemented with 1mg/ml Papain (10108014001, Sigma) for 25 min at 37° C and resuspended once during incubation. Cells were then transferred to HBSS (no calcium, no magnesium, no phenol red; 11140035, ThermoFisher Scientific) supplemented with 5%FBS, Rock inhibitor (Y-27632, Stem Cell Technologies) and 1X non-essential amino acids (11140035, ThermoFisher Scientific). Cells were further disaggregated by pipetting and filtered several times using 20 μm strainers (130-101-812, Miltenyi Biotech GmbH). To access quality live/cell death, cell size and number of clumps were measured. Samples with a viability above 65% were used for single cell sequencing. During protocol optimization, qPCR was carried out to check that gene expression levels are similar in dissociated cells and the intact hindbrain.

### 10X Genomics single-cell library preparation

A suspension of 10,000 single cells was loaded onto the 10X Genomics Single Cell 3’ Chip. cDNA synthesis and library construction were performed according to the manufacturers protocol for the Chromium Single Cell 3’ v2 protocol (PN-120233, 10X Genomics). cDNA amplification involved 12 PCR cycles. Samples were sequenced on Illumina HiSeq 4000 using 100 bp paired-end runs.

### Bioinformatic analysis of scRNA-seq data

The 10X Cell Ranger software was used to de-multiplex Illumina BCL output, create fastq files and generate single cell feature counts for each library using a transcriptome built from the zebrafish Ensembl release 89, GRCz10.

### Seurat unsupervised analysis of aggregated data

Three 10X libraries representing the 16 hpf, 24 hpf and 44 hpf stages of embryonic development were aggregated using the 10X software “cellranger aggr” function, which sub-samples reads such that all libraries have the same effective sequencing depth. Aggregated count data were further analysed using the Seurat v2.3.4 (Butler et al., 2018b) package within R v3.5.1. Lowly expressed genes with a count of >=1 in fewer than 30 cells (−0. 33%) of the data were discarded. Cell quality was assessed using some simple QC metrics and outlier cells with: (i) unique gene counts >4000 or <200, (ii) nUMI >2500 or (iii) unusually high mitochondrial RNA content (>0.05%) were removed from further analysis. Data were normalised across cells using the “LogNormalize” function with a scale factor of 10,000. A set of genes highly variable across cells was identified using the “FindVariableGenes” function (x.low.cutoff = 0.02, x.high.cutoff = 3, y.cutoff = 0.5).

Cells were scored to determine which stage of the cell cycle they were in using Seurat’s “CellCycleScoring” function and a list of cell-cycle markers from (Tirosh et al., 2016). PCA analysis on cell cycle genes revealed that cells separated entirely by phase. Given that cells were likely a mixed population of possibly quiescent stem cells and proliferating differentiated cells it was decided to regress out just the difference in G2M and S phase scores (“CC.difference”). Thus, signals separating non-cycling cells and cycling cells were maintained but differences in cell cycle phase amongst proliferating cells were removed.

Data were scaled using the “ScaleData” function, regressing out cell-cell variation caused by cell-cycle, the number of detected molecules per cell and percent mitochondrial effects (vars. to. regress = c(“CC. Difference”, “nUMI”, “percent. mito”)).

PCA analysis was performed on the scaled data using the variant genes. Significant principle components were identified by manual inspection of the top loading genes and by plotting the standard deviations of the top 100 components. Graph-based clustering of cells was performed on principle components 1-22 using Seurat’s FindClusters function (resolution = 0.6). t-SNE dimensionality reduction was run on principle components 1-22 using the “RunTSNE” function with otherwise default settings. Graphing of the output enabled visualization of cell cluster identity and marker gene expression.

Visual inspection of hindbrain and non-hindbrain marker genes suggested some clusters were constituted by contaminant non-hindbrain cells; see Supplementary Table 1 for a list of valid hindbrain cells. A new iteration of the analysis was then performed as above, this time excluding contaminant cells from the aggregated data prior to normalisation, variable gene selection, data scaling and dimension reduction (PC1-23) and cluster identification (resolution = 0.7).

Biomarkers of each cluster were identified using Wilcoxon rank sum tests using Seurat’s “FindAllMarkers” function. It was stipulated that genes must be present in 25% of the cells in a cluster and show a logFC of at least 0.25 to be considered for testing. Only positive markers were reported. The expression profile of top markers ranked by average logFC were visualised as heatmaps and dotplots of the scaled data. Cluster identity was determined using visual inspection focusing on the expression of known marker genes.

### Seurat unsupervised analysis of individual stages

Count data for individual stages were loaded directly into Seurat from the 10X results files separately, without aggregation. Downstream analysis was conducted as for the aggregated dataset. In each case the first 25 principle components were used for cluster identification (resolution = 0.6) and tSNE dimensional reduction.

### Seurat supervised clustering of ventral progenitors from individual stages

For each stage, cells identified as being ventral progenitors in the aggregate analysis were subset and subjected to supervised clustering using custom sets of marker genes to drive PCA analysis, cluster identification and tSNE dimensional reduction. For 24 hpf ventral progenitor cells, the genes used were *rfng* (boundary), *etv5b* (segment centre) and *neurog1, neurod4* (neuronal differentiation). For 44 hpf ventral progenitor cells, the list was restricted to *etv5b, neurog1* and *neurod4*.

### Pseudotime analysis of aggregated dataset using Monocle

Pseudotime analysis was conducted using the Bioconductor package Monocle v2.10.1 (Trapnell et al., 2014a) starting with the pre-filtered Seurat object. Estimates of cell size factors and dispersions were calculated using Monocle’s “estimateSizeFactors” and “estimateDispersions” functions respectively, with default settings.

Data were reduced to 2 dimensions via the Discriminative Dimensionality Reduction with Trees (DDRTree) algorithm using cluster biomarkers identified from the earlier Seurat analysis, FDR<=0.01. This created a branched pseudotime trajectory with discrete cell states on which cells were ordered. The root of the trajectory was defined as the State containing the majority of the 16 hpf progenitor cells. Top markers of cell State were identified using Monocle’s “differentialGeneTest” function (fullModelFormulaStr = “-sm.ns(State)”), restricting the search space to genes identified as significant cluster biomarkers from the Seurat analysis. Significant markers (FDR<=0.01) that were listed as being transcription factors in the AnimalTFDB3.0 database were presented on a heatmap of expression over pseudotime. Similarly, transcript factors that significantly varied in their pseudo-temporal expression pattern were also identified in a separate test (fullModelFormulaStr = “~sm.ns(Pseudotime)”).

### GENIE3 inference of regulatory networks

The Bioconductor package GENIE3 v1.4.3 (Huynh-Thu et al., 2010) was used to infer regulatory networks of genes within cells of individual developmental stages. For each stage, an expression matrix of raw gene counts, with non-hindbrain cells removed, was constructed and passed to the GENIE3 function together with a list of zebrafish transcription factors identified in the AnimalTFB3.0 database (targets = NULL, treeMethod = “RF”, K = “sqrt”, nTrees = 1000) in order to create a weighted adjacency matrix. The weights describe the likelihood of a regulator-gene / target-gene link being genuine. This matrix was converted to a table of regulatory links (regulator-gene, target-gene, link-weight). Regulator/target links with weights > 0.025 (data available in Suppl. Table 5) were visualised as an interaction directed network within Cytoscape (Shannon et al., 2003).

## ACKNOWLEDGEMENTS

We thank Andreas Sagner and Julien Delile for advice, and Qiling Xu for comments on the manuscript. We are also grateful to the Francis Crick Institute Advanced Sequencing platform and Aquatics facility for their excellent support. This work was supported by the Francis Crick Institute which receives its core funding from Cancer Research UK (FC001Z17), the UK (FC001217), the UK Medical Research Council (FC001217), and the Wellcome Trust (FC001217).

## COMPETING INTERESTS

The authors declare that they have no competing interests.

## AUTHOR CONTRIBUTIONS

Conceptualisation: M.T., D.G.W; Methodology: M.T., R.M., D.G.W.; Software: R.M.; Formal Analysis: R.M.; Investigation: M.T.; Writing - original draft: M.T., D.G.W.; Writing - review and editing: M.T., R.M., D.G.W.; Supervision: D.G.W.; Funding acquisition: D.G.W.

## SUPPLEMENTARY FIGURE LEGENDS

**Supplementary Figure 1. Quality Control matrix.**

Distribution of the number of genes per cell (nGene), the number of Unique Molecular Identifiers (UMIs) and percentage of mitochondrial reads are shown for 16 hpf (A), 24 hpf (B), 44 hpf (C) and the aggregated data set (D).

**Supplementary Figure 2. Tissue composition of hindbrain and surrounding tissues.**

tSNE representation of the three developmental stages (A-C) and the aggregated data set (D). The clustering of cells depicts their transcriptional similarity. (A’-D’) tSNE plots showing the log normalised counts of known marker genes for the different cell populations (six1 = otic vesicle; neurod1 = cranial ganglia; twist1a = neural crest; colec12 = head mesenchyme; sox3 = neural progenitors; elavl4 = neurons). Colour intensity is proportional to the expression level of a given gene. (A”-D”) tSNE map coloured based on assigned cell identity (hindbrain cells in pink, non-hindbrain cells in cyan).

**Supplementary Figure 3. Selected expression patterns of progenitors and differentiating factors at 44hpf.**

Progenitor marker *sox3* (A) and *plp1a* (B) show similar expression in the ventricular zone. *her4.4* (C), *rtca* (D), *dusp1* (E), *zbtb18* (F) and *fstl1b* (G) are present in differentiating cells showing a pattern similar to *neurog1* (F) and *neurod4* (G). *ebf2* (J) has an expression domain resembling *neurod4,* plus is also expressed in some differentiated neurons. *atp1a1b* (K) is present in glial cells. *ascl1a* (L) and *ptf1a* (M) are proneural genes found medially in differentiating cells. *prdm8* (N) has a complex expression with a medial and neurogenic domain. *atoh1a* (O) is expressed dorsally. For each gene the tSNE plot shows gene expression from 44 hpf scRNA-seq data. In situ hybridization images are shown for dorsal view, side view and transverse section at the level of r4-r5/r5-r6. scRNA-seq and in situ hybridization expression patterns strongly correlate.

**Supplementary Figure 4. Proliferation gene expression at different stages.**

For each gene, the tSNE plot shows gene expression from 16 hpf (A-D), 24 hpf (E-H) and 44 hpf (1-L) scRNA-seq data. Whole mount in situ hybridization at 44 hpf of *mki67* (M), *nusap1* (N), *ccnd1* (O) and *cdca8* (P). Dorsal view (M-P), side view (M’-P’) and 40 μm hindbrain transverse section at the level of r4-r5/r5-r6 (M”-P”).

**Supplementary Figure 5. Fgf20a-/- bulk RNA-seq identifies metrnl and fsta as new Fgf20 targets in the hindbrain.**

(A) Examples of *etv5b* expression in dissected hindbrain for wild-type (WT) and *fgf20a-l-24* hpf embryos. Five stripes of segment centre expression occur in WT embryos, together with otic vesicle and cranial ganglia expression domains. In *fgf20a-l-* embryos only weak r3 and r5 stripes are present, while otic vesicle and cranial ganglia expression domains are unaffected. Representative whole embryos are also shown. Strong expression in domains outside the hindbrain probably masks changes in the hindbrain (e.g. *etv5b).* (B) Heatmap showing RNA-seq expression levels of significantly differentially expressed genes between 4 WT and 3 *fgf20a-/*-dissected tissues. Hierarchical clustering groups the WT tissues and the mutants in separate clusters; suggesting genome wide similarities in dissected samples of the same genotype. Colour scale depicts low to high expression in blue to red shades, respectively. (C) Volcano plot shows 377 significantly downregulated genes in blue and 242 upregulated in red. *metrnl* and *fsta* are among the downregulated factors. Grey dots are non-significant genes, x-axis Log2(Fold Change) and y-axis −Log10(pvalue).

**Supplementary Figure 6. Constitutive activation of FgfR1 ectopically induces *etv5b*, *metrnl* and *fsta* expression.**

Heatmap shows RNA-seq expression levels of significantly differentially expressed genes between 4 heat shocked controls (HspCnt) and 4 heat shocked constitutive active FgfR1 (HspFgfR1CA) dissected tissues. Hierarchical clustering groups the 4 HspCnt tissues and the 4 HspFgfR1CA in separate clusters; suggesting genome wide similarities in dissected samples of the same genotype. Colours scale depicts low to high expression in blue to red shades, respectively. 8 genes are significantly downregulated in HspFgfR1CA, while 36 are upregulated. Among the upregulated genes, known Fgf signaling targets are found (e.g. *spry2, spry4* and *etv5b)* and in addition *metrnl* and *fsta* are found that are expressed in hindbrain segment centres.

## SUPPLEMENTARY FILES

**Supplementary File 1. top30_markers_per_cluster_significant_heatmap_16hpf**

Full heatmap of the top 30 significant markers per cluster at 16 hpf.

**Supplementary File 2. top30_markers_per_cluster_significant_heatmap_24hpf**

Full heatmap of the top 30 significant markers per cluster at 24 hpf.

**Supplementary File 3. top30_markers_per_cluster_significant_heatmap_44hpf**

Full heatmap of the top 30 significant markers per cluster at 44 hpf.

**Supplementary File 4. top30_markers_per_cluster_significant_heatmap_aggregate**

Full heatmap of the top 30 significant markers per cluster of the aggregate data-set.

**Supplementary File 5. top50_markers_per_cluster_ heatmap_24hpf_VP**

Full heatmap of the top 50 markers per cluster, when available, of the supervised clustering analysis done on 24 hpf Ventral Progenitors (VP).

**Supplementary File 6. TF_markers_of_state_pseudotime_heatmap**

Full heatmap of the significant TFs markers of pseudotime states.

**Supplementary File 7. TF_changing_with_pseudotime_heatmap**

Full heatmap of the significant TFs changing with pseudotime.

## SUPPLEMENTARY TABLES

**Supplementary Table 1. Hindbrain_cells**

Spreadsheet containing the names of cells considered to be hindbrain cells on the basis of marker gene expression.

**Supplementary Table 2. Gene_expression_annotation**

<u>Spreadsheet 1</u> - Table 2.1. Expression pattern summary of selected genes differentially expressed at 16 hpf.

<u>Spreadsheet 2</u> - Table 2.2. Expression pattern summary of selected genes differentially expressed at 24 hpf.

<u>Spreadsheet 3</u> - Table 2.3. Expression pattern summary of selected genes differentially expressed at 44 hpf.

<u>Spreadsheet 4</u> - Table 2.4. Expression pattern summary of selected genes differentially expressed in the Aggregate data set.

<u>Spreadsheet 5</u> - Table 2.5. Expression pattern summary of differentially expressed genes between boundary and centre progenitors at 24 hpf.

<u>Spreadsheet 6</u> - Table 2.6. Expression pattern summary of differentially expressed genes between boundary and centre progenitors at 44 hpf.

References are listed below.

**Supplementary Table 3. Fgf20aHOM_RNAseq_differential_expression_gene_significant**

Bulk RNA-seq analysis of 4 wild-type (WT) and 3 Fgf20a-/- dissected hindbrain tissues.

**Supplementary Table 4. HspFgfR1CA_RNAseq_differential_expression_gene_significant**

Bulk RNA-seq analysis of 4 heat shock control (HspCnt) and 4 heat shock constitutive active FgfR1 (HspCAFgfR1) dissected hindbrain tissues.

**Supplementary Table 5. 44hpf_GENIE3_IM0.025**

Genie3 table of interactions presenting regulatoryGene, targetGene and weight of the interaction (IM>=0.025).

